# MNetClass: A Control-Free Microbial Network Clustering Framework for Identifying Central Subcommunities Across Ecological Niches

**DOI:** 10.1101/2025.03.07.642127

**Authors:** Yihua Wang, Qingzhen Hou, Fulan Wei, Bingqiang Liu, Qiang Feng

## Abstract

Investigating microbiome subnetworks and identifying central microbes in specific ecological niches is a critical issue in human microbiome studies. Traditional methods typically require control samples, limiting the ability to study microbiomes at distinct body sites without matched controls. Moreover, some clustering methods are not well-suited for microbial data and fail to identify central subcommunities across ecological niches after clustering.

In this study, we present MNetClass, a novel microbial network clustering analysis framework. It utilizes a random walk algorithm and a rank-sum ratio-entropy weight evaluation model to classify key subnetworks and identify central microbes at any body site without the need for control samples. By applying MNetClass to microbiome data from five distinct oral sites, we successfully uncovered site-specific microbial subgroups and their central microbes. Additionally, simulations and Autism Spectrum Disorder (ASD) cohort datasets demonstrated that MNetClass outperforms existing unsupervised microbial clustering methods in terms of both accuracy and predictive power. In case studies, MNetClass identified age-related microbial communities across different oral sites, highlighting its broad applicability in microbiome research.

**IMPORTANCE:** MNetClass provides a valuable tool for microbiome network analysis, enabling the identification of key microbial subcommunities across diverse ecological niches. Implemented as an R package (https://github.com/YihuaWWW/MNetClass), it offers broad accessibility for researchers. Here, we systematically benchmarked MNetClass against existing microbial clustering methods across multiple datasets using various performance metrics, demonstrating its superior efficacy. Notably, MNetClass operates without the need for control groups and effectively identifies central microbes, highlighting its potential as a robust framework for advancing microbiome research.

## INTRODUCTION

Microbes are widely distributed in the human body, and together with our tissues and organs, they form an organic component of the human body, playing an important role in maintaining human health^1,2^. Due to the diverse physiological characteristics and microenvironments, the composition of microorganisms differs in each body site, with some differences being significant^3^. Similarly, microbes have preferences for body sites, and ectopic colonization of certain microbes is a risk factor for many major diseases^4^. For example, *Fusobacterium nucleatum* is a common bacterium in the oral cavity and is an important pathogenic bacterium of colon cancer after it colonizes the intestine^5^. *Porphyromonas gingivalis* is another prevalent oral bacterium that can induce periodontitis in the oral cavity and could serve as a promoter of esophageal cancer when it colonizes the esophagus^6^. Therefore, illuminating the composition and ecological characteristics of microbes in each body site is of great significance for understanding the relationships between human health and symbiotic microbes.

Currently, identifying microbes closely related to specific human diseases is an important discussion in the study of disease causality^7^. Previous studies have mainly relied on statistical methods, which help to identify microbes with significant differences by comparing microbial abundance across different groups of study cohorts and control groups. However, no microbe exists in isolation; each is intricately linked with others and forms a well-balanced ecological network^8^, which creates better living conditions through mutual symbiosis and cooperative metabolism or enables the microbes to evade host immune surveillance^9^. The microbiome-wide association study (MWAS)^10^ primarily employs differential abundance analysis of each single microbe to identify characteristic microbes that are significantly differently distributed in disease groups. This ignores the interactions among microbes, resulting in a relatively low alignment with actual inter-microbial relationships within the ecological network.

To address this issue, multiple network-based approaches have been developed for exploring the composition and interactions within the microbial ecosystem^11,12^. For example, NetShift^13^ introduces a neighbor shift score (NESH) to analyze the variability of node neighbors in the network, incorporating node betweenness to quantify the driving force of species in the network and thus identifying the “drivers” of microbial network change. NetMoss^14^ is used to evaluate the changes in microbial network modules to identify biomarkers related to various diseases, exhibiting superior performance in removing batch effects during data integration. Manta^15^ is a network clustering algorithm that leverages negative edges and distinguishes between weak and strong cluster assignments, offering superior robustness to noise and better performance in identifying biologically relevant groups. However, most of these network analysis methods either require control groups or fail to accurately identify central subcommunities within the microbial interaction network after clustering.

In this paper, we develop a microbial network clustering framework named MNetClass. By accounting for interactions between microbes, MNetClass can classify unique subgroups for each location while also exploring the network’s characteristic microbes and their intermicrobial relationships using a Walktrap^16-19^ algorithm and rank-sum ratio-entropy weight evaluation model^14,20^. Unlike other methods, MNetClass operates without the need for control samples and directly analyzes microbial features based on their abundance and interactions within the ecological niche. After network partitioning, it effectively identifies central subcommunities, providing deeper insights into microbiome organization and structure. We demonstrate the capabilities and versatility of MNetClass on both simulated and real microbiome datasets, where it outperforms seven other unsupervised microbial clustering algorithms. MNetClass reliably identifies site-specific microbial communities in oral datasets and shows superior predictive performance in cross-cohort Autism Spectrum Disorder (ASD) data. In case study, it can classify age-related microbial communities across oral sites. MNetClass is implemented in an open-source R package and is freely available at https://github.com/YihuaWWW/MNetClass, providing a novel all-in-one network analysis service for microbiome research.

## MATERIALS AND METHODS

### Overview of MNetClass

In general, our MNetClass framework mainly contains 3 steps: (1) constructing a microbial association network from a microbial abundance matrix; (2) partitioning the network into subnetworks using a random walk algorithm; and (3) evaluating subnetworks and network nodes (microbes) based on topological properties, followed by scoring and ranking them through an integrated rank-sum ratio–entropy weight evaluation model.

For the microbial association network, we utilize the classic Spearman’s correlation method. The microbial correlation network is characterized as a weighted graph *G* = (*V, E, W*). In the network, nodes (*V*) represent bacteria, with the weight of a node denoting the average relative abundance of that bacteria in the ecological niche; and edges (*E*) represent interbacterial interactions, with their weights (*W*) being measured by Spearman’s correlation coefficient between microbial taxa. Within a network, communities are subgraphs (or subnetworks) containing more internal connections than external connections, representing microbial subpopulations with strong interaction relationships in the microbial ecosystem^21^.

Next, a community partitioning algorithm based on random walks is employed to explore specific, highly interactive microbiota subpopulations within a particular microbial ecosystem. For specificity, scoring models are utilized to compare the topological properties of subnetworks, including network local properties such as degree centrality^22^, betweenness centrality^23,24^, closeness centrality^23,25^, and eigenvector centrality^26,27^, as well as global network properties, such as graph diameter^28^, modularity^29-31^, density^32^, and clustering coefficient^33^.

To mitigate the influences of magnitude differences among various indicators on the assessment of network topological properties, the integrated rank-sum ratio– entropy weight evaluation model is leveraged to holistically evaluate the composite topological properties of the microbial network in a given niche. The subnetworks or nodes (microbes) with the highest composite score are then selected as the characteristic subgroup and network node bacteria for that particular niche.

#### 1. Data Pre-processing: Normalization and Filtration of Low-Abundance, Low-Frequency Microorganisms

Microbiome data processing begins with a nonnegative matrix containing the abundant data of all annotated microbial species in each sample. The data include the total read counts 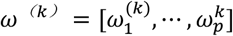 for *p* taxa in sample *k*. Here, 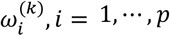 represents the read counts of taxa *i* in sample *k*. Due to variations in sequencing depth across samples and the influence of sequencing technologies, the abundance matrix is often sparse, containing a large proportion of zeros. Therefore, we performed preprocessing on the data.

First, to make the read counts comparable across samples, the microbial sequence counts are normalized into relative abundance using the total sum scaling (TSS) normalization technique^34,35^ (step a in Figure 1). However, proportions are typically compositional, even if the raw data are not. Aitchison recommended the application of the centered log-ratio (clr) transformation to convert compositional data from the simplex to real space^34^. Then, the taxa with low abundance or low frequency (step b in Figure 1) are filtered out to reduce the sparsity of the relative abundance matrix.

**Figure 1.**
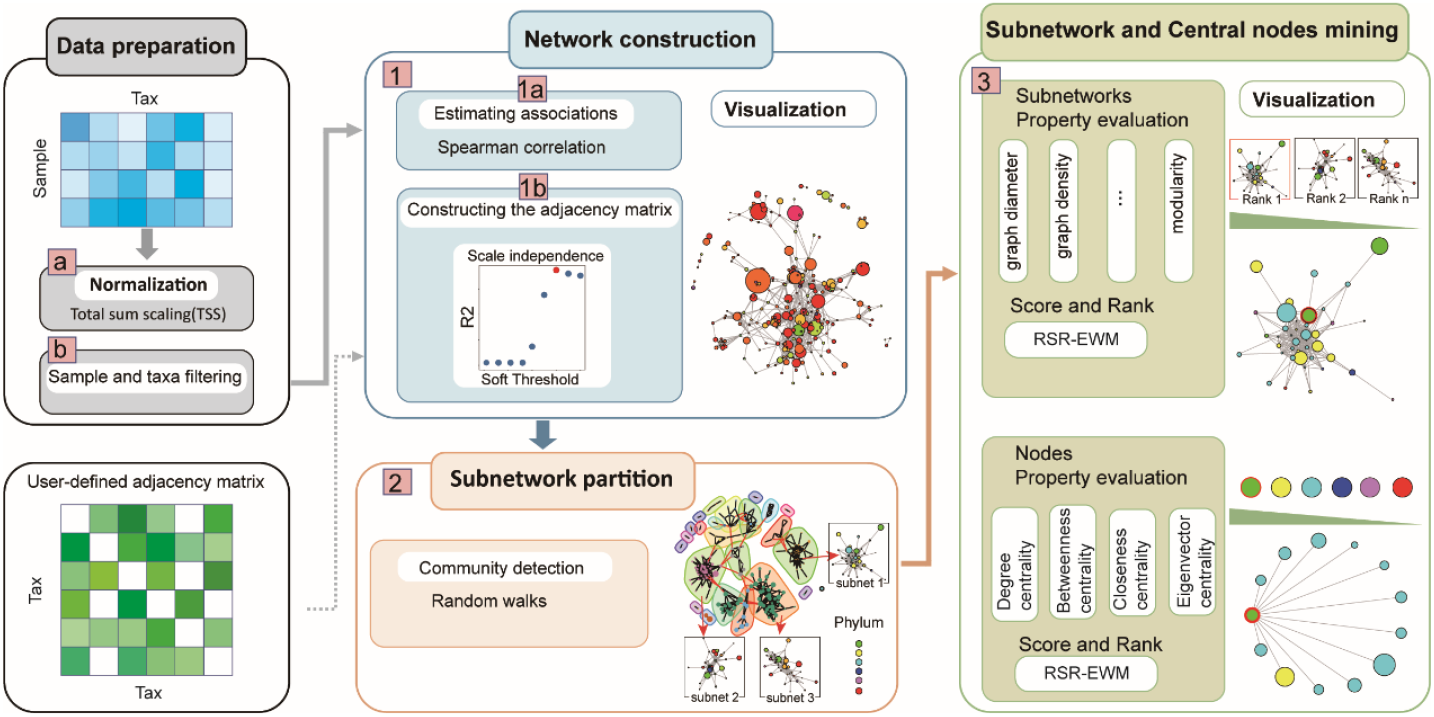
The simplified diagram of MNetClass’s analysis pipeline. Data pre-processing to normalize and filter low-abundance & low-frequency microorganisms. 1. Constructing a microbial association network. 2. Partitioning the network into subnetworks. 3. Evaluating and scoring microbial topological properties of subnetworks and central node microbes

#### 2. Constructing a microbial association network

##### 2.1 Correlation analysis among all microbes in a given niche

To ensure wide applicability of MNetClass, the classical Spearman’s rank correlation coefficient^36,37^ is used to measure interactions between microbial taxa (step 1a in Figure 1) to obtain the association matrix for microbial interactions in a given niche. Correlation-based techniques remain one of the most popular approaches for analyzing microbial networks in human gut, oral, and soil microbiomes due to their low computational complexity, simplicity, flexibility, and ease of operation^38,39^. These methods have led to significant research discoveries in understanding microbial interactions^40-43^.

For a given set of n samples with paired relative abundance data for taxa *X* and taxa *Y*, {(*x*_1_, *y*_1_), (*x*_2_, *y*_2_), ⋯, (*x*_*n*_, *y*_*n*_)}, the Spearman correlation coefficient *r*_*s*_ is defined as^44^ 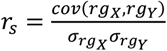, where *rg*_*x*_ and *rg*_*y*_ represent the ranks of the relative abundance data for taxa *X* and *Y*, respectively. cov(∗,∗) denotes the covariance of the rank variables, and 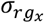 and 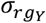 represent the standard deviations of the rank variables *rg*_*X*_ and *rg*_*Y*_.

##### 2.2 Construction of the Microbial Adjacency Matrix

Since the estimated correlations are generally nonzero, directly using them as the adjacency matrix results in a densely connected network. Therefore, the matrix A of the Spearman correlation coefficient is processed through two sparsification steps. First, we introduce a defining threshold and keep the pairs of microbes with absolute correlation coefficients above this threshold^24,45^. Second, Student’s t test^46^ is employed to identify correlations that are significantly different from zero. Then, we generate a sparse adjacency matrix *A*^*p*×*p*^ (step 1b of Figure 1). In this matrix, the entries *a*_*ij*_ represent the correlation coefficients between species *i* and *j*, and these values serve as the weights for the edges of the microbial network, with the nodes representing the microbial taxa. The diagonal elements, *a*_*ii*_, are set to zero, as they represent self-correlations, which are not relevant in this context.

We selected a biologically motivated approach suitable for scale-free networks to determine the optimal correlation coefficient threshold, rather than relying on an entirely arbitrary choice. Scale-free networks exhibit a degree distribution that follows a power-law pattern, characterized by a small number of highly connected nodes (hubs) and a large number of sparsely connected nodes. This distribution has been observed in various biological networks, including microbial co-occurrence networks ^43,47-50^. It is also commonly used as a threshold criterion in the widely adopted WGCNA package ^51^.

To identify the optimal threshold that ensures the resulting network aligns with the scale-free property, we tested a range of correlation thresholds (0-1, in increments of 0.1). For each threshold, the degree distribution of the network was calculated using the igraph package^52^. We then fitted the degree distribution to a power-law model using the nonlinear regression function nls in R, based on the frequency distribution of node degrees. The fitted power-law equation was used to predict the response variable (degree frequency) as a function of the predictor variable (degree). The goodness-of-fit was evaluated by calculating *R*^2^, and the significance of the *R*^2^ value was tested using a permutation test to obtain the associated p-value^53^. Finally, the threshold corresponding to the maximum *R*^2^ value with *p* ≤ 0.05 was selected as the optimal correlation coefficient threshold.

#### 3. Partitioning the network into subnetworks

Previous studies have demonstrated that subnetworks frequently identify functionally coherent microbial groups^54,55^. Community detection methods are instrumental in identifying subgroups within microbial communities that exhibit strong associations. The Walktrap algorithm^18,19,56^ is employed to partition the sparse weighted network into multiple relatively independent subnetworks. For an undirected network *G* = (*V, E, W*) consisting of *n* = |*V*| nodes and *M* = |*E*| edges, this algorithm can calculate a community structure with a time complexity of *O*(*MnH*), where *H* is the height of the corresponding dendrogram. The primary advantage of this algorithm is that it can produce robust results rapidly without requiring the number and size of the communities as parameters^17^. Walktrap algorithm is based on the general idea that, if one performs random walks on a network, then the walks are more likely to stay within the same cluster due to the assumed higher level of interconnectedness within clusters. Walktrap is slightly slower than fast greedy approach, but it is shown to be more accurate^18^. Other algorithms for community detection can also be used in this method; these algorithms are presented in Supplementary Material 1.

The measure of similarity between nodes is based on the distance values obtained from the random walk algorithm. Walktrap utilizes certain characteristics of random walks on graphs to define a measure of structural similarity between nodes and communities, thereby defining a distance *r*^18,57^, as shown in Equations 1 and 2. Here, the distance is used in hierarchical clustering algorithms to merge nodes (microbes) into communities.

**Definition 1** Let *i* and *j* be two nodes in the graph,

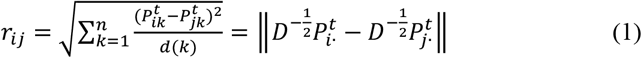

where ‖·‖ denotes the Euclidean norm on ℝ^n^. r_ij_ represents the distance between nodes *i* and *j*. Let 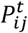 represent the probability of a random walk of length *t* from node *i* to node *j*. Furthermore, *P* denotes the transition matrix of the random walk process. Specifically, *P* is defined as *P* = *D*^−1^*A*, where *A* is the adjacency matrix of graph *G*, and *D* is the degree matrix (∀*i, D*_*ii*_ = *d*(*i*) and *D*_*ij*_ = 0 for *i* ≠ *j*).

**Definition 2** Let *C*_1_, *C*_2_ ⊂ *V* be two communities of the graph. We define the distance between the two communities, denoted as *r*_*C*1_*C*_2_, as

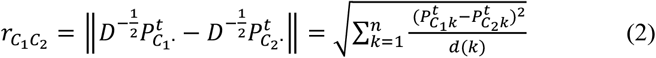

Here, 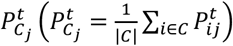 represents the probability of reaching node *j* from community *C* within *t* steps. 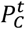 is a probability vector. *D* denotes the degree matrix.

#### 4. Evaluating and scoring microbial topological properties of subnetworks and central node microbes

##### 4.1 Evaluating and scoring microbial subnetworks

In microbial community network analysis, the topological properties of subnetworks can be leveraged to identify key microbial taxa. Studies have indicated that microorganisms with higher topological centrality are more likely to play pivotal roles within the community^42,58^. Therefore, we conducted a topological analysis of the subnetworks derived from community detection, calculating their respective network properties. In each niche, the different subnetworks ranked in the top k percentile by node count (k=20) are selected and evaluated by ten same network topological property indicators, including the graph diameter^28^, graph density^32^, average path length^28^, average degree^59^, edge and vertex connectivity^32^, clustering coefficient^33^, and modularity^29^; these are described in Supplementary Material 2.

Next, we identified key subnetworks based on their topological properties. According to previous studies^60-62^, selecting subnetworks using multiple topological properties is more reliable than relying on a single property, as the resulting microbial communities are more likely to represent keystone species. Inspired by this approach, we incorporated all the aforementioned topological properties as selection criteria. However, the scales and relative importance of these properties vary. To address this, an integrated rank-sum ratio–entropy weight (RSR-EWM) evaluation model^14^ is used to assess the composite topological properties of each subnetwork. The subnetwork with a topological property score of “1” is chosen to represent the characteristic subgroups in this niche. Since the topological property values used in the process are rank-ordered, representing the relative magnitudes of the data rather than the data itself, there are no special requirements for indicator selection. This makes the rank-sum ratio (RSR) applicable to various evaluation subjects and capable of identifying minor changes in the objects of study^20^. We introduce the entropy weight method (EWM) to perform weighted analysis of evaluation indicators^63^. The EWM is an objective evaluation method following the principle that the greater the dispersion of an indicator is, the lower its information entropy and the more information it contains.

The specific evaluation process of the RSR-EWM is as follows:

###### (i) Compiling the topological property evaluation data into a matrix

Assuming *M* subnetworks require comprehensive topological property assessment and *n* evaluative indicators are used, an *M* by *n* data matrix can be generated, *X* = (*X*_*ij*_)_*M*×*n*_. Here, *X*_*ij*_ represents the numerical value of the *j*th topological property indicator of the *i*th subnetwork.

###### (ii) Standardizing the subnetwork topological property metrics

For the positive network topological property, where larger values correspond to better network topological properties, the standardization process is expressed as Equation (3). For the negative indicators, the standardization process is expressed as Equation (4).

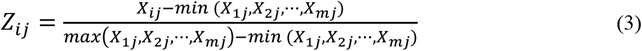

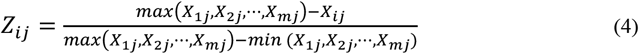

###### (iii) Computing the rank of each indicator

From the *M* by *n* data matrix *X* = (*X*_*ij*_)_*M*×*n*_, generated from the *M* subnetworks and *n* evaluative indicators, we compile the ranks for each evaluation object for every indicator. The positive network topological property indicators are ranked in ascending order. When identical data are observed for the same indicator, an average rank is assigned, resulting in the rank matrix, *R*.

###### (iv) Weighting of each indicator by EWM

**Step 1:** Calculation of the proportion, *p*_*ij*_, that the *i*th evaluation object holds for the *j* th indicator. This proportion, *p*_*ij*_, is utilized as the probability in the calculation of relative entropy, as described in Equation (5).

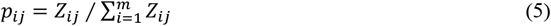

**Step 2:** Calculation of the information entropy, *e*_*j*_, for each indicator, as detailed in Equation (6).

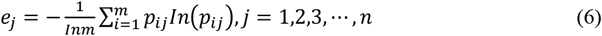

**Step 3:** Normalization of the information entropy to ascertain the entropy weight, *W*_*j*_, of each indicator, as described in Equation (7).

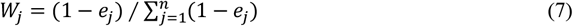

###### (v) Computing and ordering the weighted rank-sum ratio (WRSR)

Calculation of the WRSR as described in Equation (8).

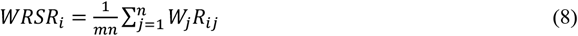

###### (vi) Converting the distribution of *WRSR*_*i*_ into probability units

The *WRSR*_*i*_ distribution is represented by the cumulative frequency in the probit unit table.

**Step 1:** Arrange the values of *WRSR*_*i*_ in ascending order.

**Step 2:** List the frequency *f* of each group and calculate the cumulative frequency ∑ *f*.

**Step 3:** Determine the average rank 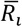 of each evaluation object. For *WRSR*_*i*_ with a frequency equal to 1, the value of 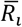 *WRSR*_*i*_ with a frequency not equal to 1, the value of 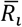 is the rank of *WRSR*_*i*_. For is the average of each rank.

**Step 4:** Calculate the downward cumulative frequency as 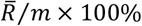 and correct the final item with (1 − 1/4*M*) × 100%.

**Step 5:** Convert the downward cumulative frequency to the *Probit* probability unit. The *Probit* is the standard normal deviation µ plus 5 corresponding to the cumulative frequencies, as referenced in the “comparison table of percentages and probability units”^64^.

###### (vii) Computing the value of 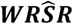

Taking the *Probit* probability unit corresponding to the cumulative frequency as the independent variable and *WRSR*_*i*_ as the dependent variable, we compute the linear regression equation as depicted in Equation (8). This regression equation is then subjected to rigorous statistical testing.

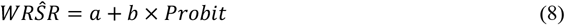

###### (viii) Ranking and Binning

In accordance with the probabilistic unit *Probit* and the adjusted rank-sum ratio value 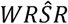,binning is conducted.

Based on the scores derived from the RSR-EWM model, the top-scoring subnetworks are selected for each respective group in the study.

##### 4.2 Evaluating and scoring central node microbes

Furthermore, within the selected subnetworks, we identified nodes with favorable composite topological properties, which we refer to as “central node microbes”. This approach is analogous to selecting hub microbes, as described in previous studies, which are characterized by having the highest degree of connectivity ^65^. These central node microbes are likely to play significant physiological roles within the selected subnetworks^60^.

The pipeline of central node microbes is similar to that of subnetwork analysis. We assess the topological attributes of the chosen subnetwork nodes (microbes) using five network centrality measures: degree^23^, weighted degree^23^, betweenness^23,24^, closeness^23^, and eigenvector centrality^26,27^, as provided in Supplementary Material 2. Next, a node’s composite score is determined using an RSR-EWM evaluation model. Nodes highly ranked by composite topological property indicators are selected as the core microbes within this characteristic subgroup^66^.

#### 5. Implementation of the MNetClass as a R package

The MNetClass methodology has been implemented as an R package for assisting easy analysis of microbial association networks obtained from distinct body sites of human microbiome studies available at https://github.com/YihuaWWW/MNetClass. The input data for this R package consist of two components: a microbial abundance and the corresponding “species-phylum” or “genus-phylum” relationship. MNetClass comprises three functions: “comm”, “netscore” and “nodescore”. The “comm” function enables the construction of correlation networks, partition of networks, calculation of network and node topological properties, as well as visualization of association networks and network partitioning results. The “netscore” and “nodescore” functions, respectively, facilitate the comprehensive scoring and ranking of subnetworks and node topological properties using RSR-EWM.

### 6. Datasets

#### 16S rRNA sequencing data from distinct oral sites

We collected samples from the gingival crevicular fluid (GCF), dental plaque (P), buccal mucosa (B), saliva (S), and tongue coating (T), of 21 volunteers and all samples were analyzed by 16S rRNA sequencing. After primary sequencing data quality control and clean read clustering, we annotated the operational taxonomic units (OTUs) by the Ribosomal Database Project (RDP version 11.5) and obtained a total of 420 species, 517 genera, and 31 phyla, as described in Supplementary Material 3 and Supplementary Table 1.

**Table 1.** EWM of topological property evaluation of each subnetwork.

|  | Vertex | Edge | Graph | Graph | Average | Edges | Nodes | Clustering | Modularity | Average |
| --- | --- | --- | --- | --- | --- | --- | --- | --- | --- | --- |
|  | number | number | diameter | density | degree | connectivity | connectivity | coefficient |  | path |
|  |  |  |  |  |  |  |  |  |  | length |
| $e_j$ | 0.780 | 0.654 | 0.525 | 0.825 | 0.732 | 0.723 | 0.723 | 0.905 | 0.642 | 0.894 |
| $W_j$ | 0.085 | 0.133 | 0.183 | 0.067 | 0.103 | 0.107 | 0.107 | 0.037 | 0.138 | 0.041 |

#### Synthetic data sets

We utilized a modified version of the FABIA R package^67^ to simulate microbial abundance matrices, which is designed for generating synthetic data to evaluate biclustering algorithms. The makeFabiaDatablocksPos function was adapted to generate biclusters at predefined locations, which were designated as true-positive clusters. The simulation was repeated 50 times to produce 50 sets of synthetic microbial abundance data. The constructed matrices were subsequently used to infer correlation networks.

#### Gut microbiome data across Autism Spectrum Disorder cohorts

In this study, we analyzed gut microbiome data and metadata from individuals with Autism Spectrum Disorder (ASD) sourced from publicly available datasets described in Xu et al. (2022)^68^. The data comprise 1019 samples (569ASD and 450 healthy controls) from 10 cohorts, with detailed metadata including age, country, sequencing region and platform. Sequencing was performed using 16S rRNA gene sequencing, and taxonomic profiling was conducted using the RDP taxonomy outline (version 14). To ensure comparability across cohorts, sequences were normalized to relative abundances. This dataset provides a robust resource for studying gut microbial patterns in ASD and enables cross-cohort comparisons to assess the consistency of microbial signatures.

#### Oral microbiome data stratified by age

In this study, we utilized oral microbiome datasets from the work of Liu et al. (2020)^69^, which examined microbial community changes across five age groups (age 11-15, 18-20, 28-32, 38-45, 50–65) and three oral sites of healthy people (gingival crevicular fluid (GCF), saliva (SAL) and tongue back (TB)). The dataset includes 179 samples, with detailed metadata describing participant age, gender and sampling sites. Sequencing was performed using 16S rRNA gene sequencing on the Illumina MiSeq platform, targeting the V3–V4 region. The original study revealed significant shifts in microbial composition both with increasing age and across different oral sites. Our study leveraged this dataset to show the real-world applicability of MNetClass and performed case analyses.

## RESULTS

### Application of MNetClass framework on 16S rRNA sequencing data from distinct oral sites

To test the efficacy of MNetClass, we implemented our pipeline on collected samples from the gingival crevicular fluid (GCF), dental plaque (P), buccal mucosa (B), saliva (S), and tongue coating (T) of 21 volunteers.

#### 1. Data Pre-processing: Data filtering, normalization, and zero handling

362, 401, 385, 346, and 345 species were obtained in the oral site GCF, P, B, S, and T, respectively. We normalized the microbial species counts into relative abundance at each oral site and filtered out the species with a relative abundance <0.0002 or an occurrence < 30%.

#### 2. Construction of microbial association networks

For the selection of the Spearman correlation coefficient threshold, we constructed microbial association networks for five different oral sites at various thresholds, and calculated the *R*^2^ and P-values for the fit to a scale-free network (Figure 2(a)). We found that, for all five sites, the network constructed with a threshold of 0.8 yielded the highest *R*^2^, indicating the best fit to a scale-free network. Therefore, we set the Spearman correlation coefficient threshold to 0.8 for network construction using the microbial datasets from these oral sites.

**Figure 2.**
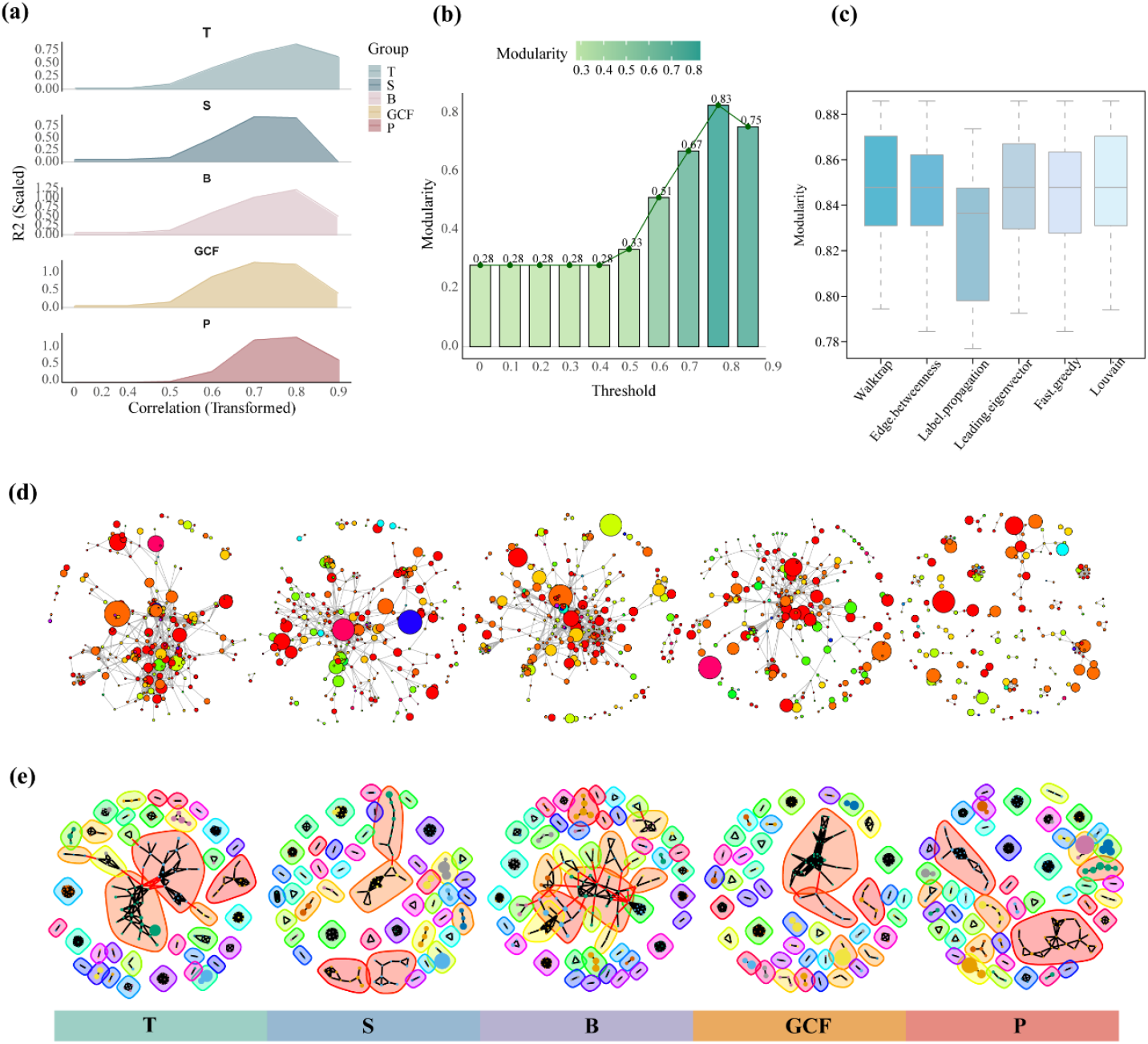
Construction and partitioning of correlation networks. (a) Distribution of *R*^2^ values for the scale-free network fit at different correlation coefficient thresholds. Different colors represent five oral sites: gingival crevicular fluid (GCF), dental plaque (P), buccal mucosa (B), saliva (S), and tongue coating (T). (b) Modularity of subnetworks after partitioning based Walktrap at different Spearman coefficient thresholds. (c) Modularity of subnetworks after partitioning using six community detection algorithms. (d) Visualization of correlation networks constructed for the five oral sites. Nodes represent bacteria, with node size corresponding to relative bacterial abundance. Node color indicates distinct phylum, and edges between nodes signify interactions between bacteria. (e) Visualization of the partitioned subnetworks. Node color represents the bacteria belonging to different clusters. Black edges indicate positively weighted interactions, signifying positive correlations between the connected bacteria, while red edges represent negatively weighted interactions, indicating negative correlations.

Additionally, using the dorsum of the tongue data as an example, we tested the modularity of subnetworks formed at different correlation coefficient thresholds (Figure 2(b)). We observed that a threshold of 0.8 resulted in higher modularity, facilitating better network partitioning.

Spearman correlation coefficients were applied and filtered at each oral site with a correlation coefficient > 0.8 and corrected P value < 0.05. The retained pairs of species were used to construct the adjacency matrices. For each oral site, “GCF”, “P”, “B”, “S”, and “T” obtained 2360, 2577, 3179, 2470, and 3333 pairs of correlated species, respectively, as described in Supplementary Table 2.

**Table 2.**
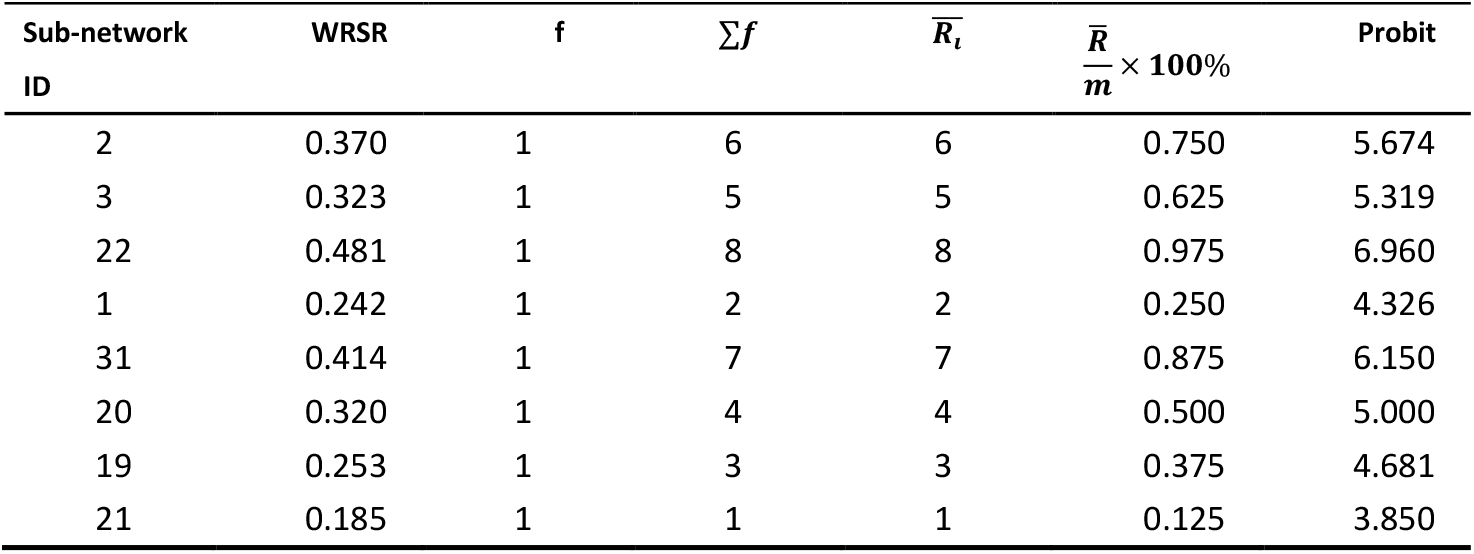
Distribution condition of statistical WRSR.

| Sub-network<br>ID | WRSR | f | $\Sigma f$ | $\overline{R}_i$ | $\frac{\overline{R}}{m} \times 100\%$ | Probit |
| --- | --- | --- | --- | --- | --- | --- |
| 2 | 0.370 | 1 | 6 | 6 | 0.750 | 5.674 |
| 3 | 0.323 | 1 | 5 | 5 | 0.625 | 5.319 |
| 22 | 0.481 | 1 | 8 | 8 | 0.975 | 6.960 |
| 1 | 0.242 | 1 | 2 | 2 | 0.250 | 4.326 |
| 31 | 0.414 | 1 | 7 | 7 | 0.875 | 6.150 |
| 20 | 0.320 | 1 | 4 | 4 | 0.500 | 5.000 |
| 19 | 0.253 | 1 | 3 | 3 | 0.375 | 4.681 |
| 21 | 0.185 | 1 | 1 | 1 | 0.125 | 3.850 |

#### 3. Variation of network partition in five oral sites

The correlation networks for each oral site were constructed based on the adjacency matrices (Figure 2(a)). Subnetwork partitioning was then performed using the Walktrap community partition algorithm, with the random walk step length set to four. Additionally, we tested five other community detection algorithms and compared the modularity of the subnetworks they generated. As shown in Figure 2(c), the Walktrap community partition algorithm resulted in higher modularity compared to the other algorithms, although the difference was not statistically significant. The partitioning networks of five oral sites are shown in Figure 2(b) and Supplementary Table 3.

**Table 3.** Topological properties ranking results for eight subnetworks.

| Grade | Probit | $W\widehat{R}S\widehat{R}$ | Sub-network Number |
| --- | --- | --- | --- |
| Excellent | $\geq 5.5$ | $\geq 0.348$ | 22, 31, 2 |
| Moderate | 4 ~ 5.5 | 0.204 ~ 0.348 | 3, 20, 19, 1 |
| Poor | $< 4$ | $\leq 0.204$ | 21 |

Specifically, “GCF” is divided into 43 subnetworks, with three encompassing more than ten nodes (Supplementary Table 3-1). “P” is partitioned into 46 subnetworks, with four encompassing more than ten nodes (Supplementary Table 3-2). “B” is partitioned into 49 subnetworks, with four encompassing more than ten nodes (Supplementary Table 3-3). “S” is divided into 45 subnetworks, with six encompassing more than ten nodes (Supplementary Table 3-4). “T” is partitioned into 39 subnetworks, with five encompassing more than ten nodes (Supplementary Table 3-5).

#### 4. Identification of subnetworks and central node microbes

For each oral site, the top 20% of subnetworks (k=20) in terms of node count were selected and evaluated by ten kinds of network topological property indicators, which are shown in Supplementary Table 4. Next, the comprehensive topological properties of each subnetwork were evaluated, and a score-based ranking order was obtained, as shown in Supplementary Material 5. The subnetworks with the highest scores were selected as the dominant subgroups in each group, as shown in Figure 3(a) and Supplementary Figure 1-4.

**Table 4.** Information entropy and weights of the five indexes in topological property evaluation of nodes in subnetwork 2.

|  | Degree | Weighted Degree | Betweenness | Closeness | Eigenvector |
| --- | --- | --- | --- | --- | --- |
|  | Centrality | Centrality | Centrality | Centrality | Centrality |
| $e_j$ | 0.876 | 0.880 | 0.667 | 0.946 | 0.843 |
| $W_j$ | 0.158 | 0.153 | 0.423 | 0.068 | 0.199 |

**Figure 3.**
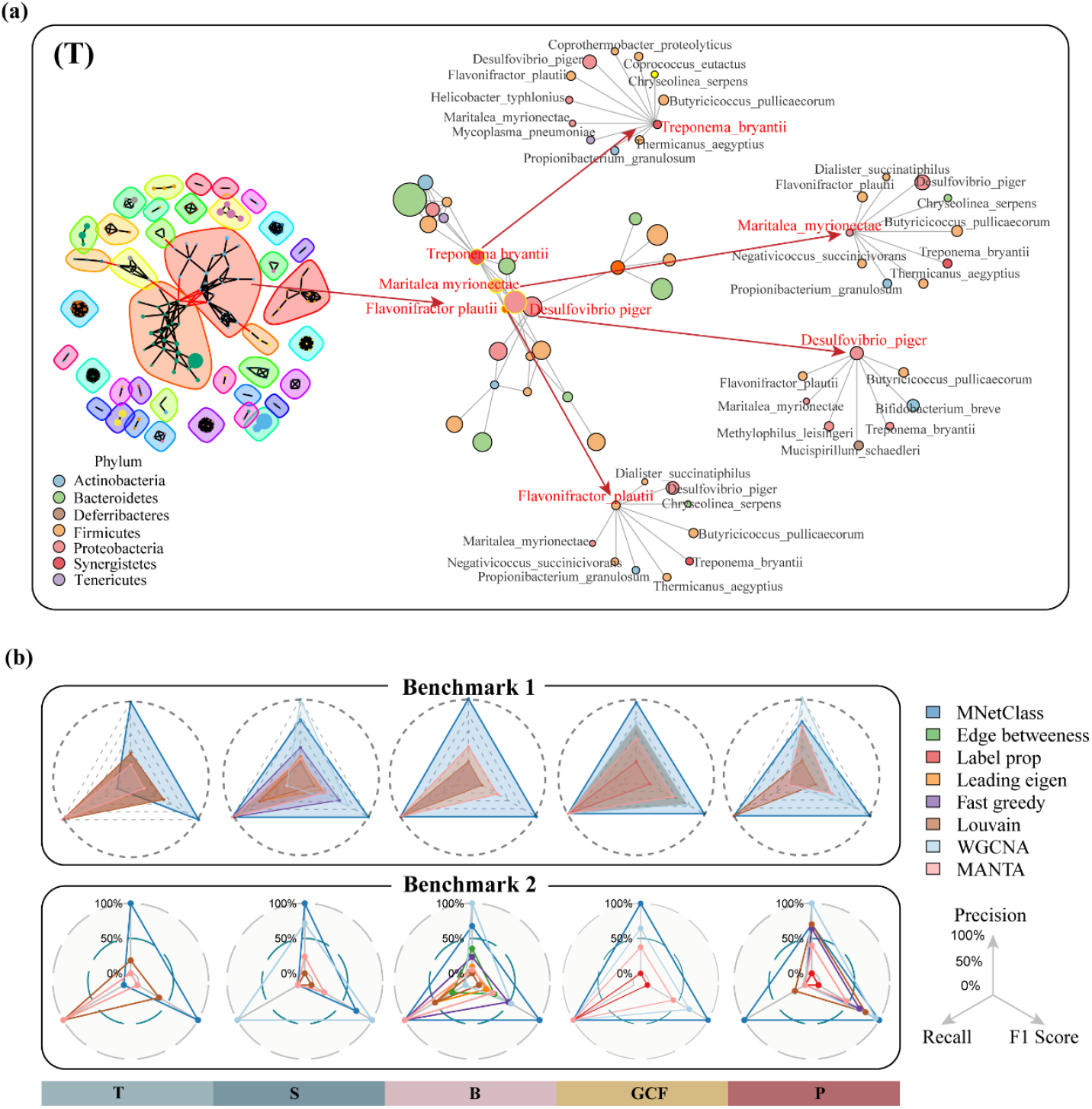
Identification and evaluation of key subnetworks and central microbes. (a) Key subnetworks and central microbes at the tongue coating (T) identified using the Walktrap algorithm and the integrated rank-sum ratio entropy weight evaluation model (RSR-EWM). The left panel shows a visualization of the subnetwork partitioning results. The middle panel highlights the important subnetworks selected from the left panel. The right panel displays the key microbial nodes and their associated bacteria selected from the middle panel. Red labels denote key microbial nodes, while black labels indicate bacteria associated with them. (b) Radar plot comparing the performance of MNetClass with seven other algorithms on two benchmarks, evaluated using Precision, Recall, and F1-score. Benchmark1 is based on EHOMD. Benchmark2 is based on meta-analysis.

Here, the analysis of subnetworks and central microbes at the niche of the tongue is detailed as an example.

##### 4.1 Evaluating and scoring of microbial subnetworks

1. The original evaluation matrix is constructed based on the scores of the ten indicators for the eight subnetworks, as shown in Supplementary Table 4-5. The matrix is presented below:

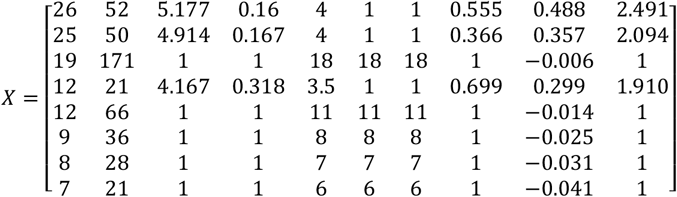
2. The subnetwork topological property metrics are standardized using Equation (1). The detailed results are presented below:

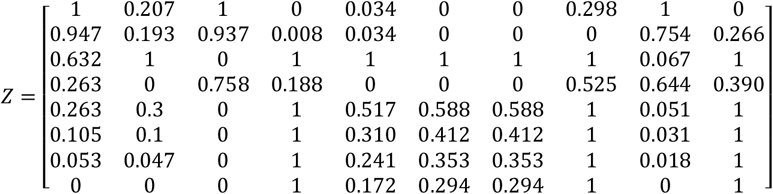
3. The Rank *R*_*ij*_ of each metric of subnetwork topological properties is calculated. Among the ten network topological property indicators, nine positive metrics are arranged in ascending order. The final indicator, “average path length,” is a negative metric and arranged in descending order. Identical values for the same metric are given average ranks, producing the rank matrix *R*. The results are provided below:

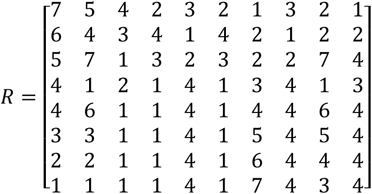
4. Weights are determined using EWM. Based on Equation (4), the information entropy *e*_*j*_ and entropy weight *W*_*j*_ for ten evaluation metrics are calculated. The results are presented in Table 1.
5. The *WRSR* values for the eight subnetworks are computed using Equation (5), with the results provided below:

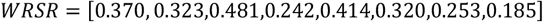
6. Distribution statistics of *WRSR* are identified. In this step, the frequency, cumulative frequency, average rank, downward cumulative rating, and probability unit are tabulated in Table 2. A higher rank indicates a better evaluation of the subject.
7. With the *Probit* values corresponding to cumulative frequencies as the independent variable and *WRSR* values as the dependent variable, the regression equation is computed using the least squares method: 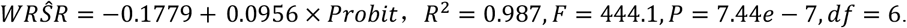.
8. The topological properties of the subnetworks are ranked based on *Probit* and 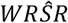,with the results presented in Table 3.

##### 4.2 Evaluating and scoring of central node microbes at Dorsal Tongue site

Similar to the evaluation of subnetwork composite topological properties, we employ an integrated rank-sum ratio–entropy weight evaluation model to evaluate the comprehensive topological attributes of all nodes within subnetworks 22, 31 and 2. This decision is based on the scores of five indicators for nodes within the excellent subnetworks 22, 31 and 2, as outlined in Supplementary Material 2. Subnetworks 22 and 31 are regular graphs, in which all nodes have the same degree, and each node has the same five topological properties, so we use the evaluation model to evaluate only the combined scores of these five topological properties for all nodes in subnetwork 2. Since the five topological properties of the nodes also serve as positive indicators, they are ranked in ascending order. Employing the same computation method, the information entropy and weight for the five topological properties of nodes are calculated. These results are provided in Table 4. The final graded comprehensive topological properties for nodes in subnetwork 2 are presented in Table 5.

**Table 5.** Topological properties ranking results for nodes of subnetwork 2.

| Grade | Probit | $WR\hat{S}R$ | Node Index |
| --- | --- | --- | --- |
| Excellent | $\geq 6$ | $\geq 1.057$ | 25, 10, 16, 13 |
| Moderate | 4 ~ 6 | 0.597 ~ 1.057 | 8, 20, 9, 5, 19, 21, 14, 11,<br>6, 3, 4, 24, 22, 7, 18, 23,<br>2, 17 |
| Poor | $< 4$ | $\leq 0.597$ | 15, 1, 12, 0 |

At site T, the Walktrap algorithm divided the correlation network into 39 subnetworks. The top eight subnetworks with the highest number of nodes (microbes) are subnetworks “2”, “3”, “22”, “1”, “31”, “20”, “19” and “21”. Then, the RSR-EWM evaluation algorithm is used to score the comprehensive topological properties of these subnetworks, with the highest ranking being “2”, which contains 26 nodes, 52 edges, a graph diameter of 5.177, a graph density of 0.160, etc. (Table 6). Next, we comprehensively scored five topological property indicators for each of the 26 nodes in subnetwork “2” and found that *Treponema bryantii, Desulfovibrio piger, Maritalea myrionectae*, and *Flavonifractor plautii* were the top four highest scoring bacteria (Table 7 and Figure 3(a)), which were the central bacteria in subnetwork “2”. Using the same pipeline, the central bacteria of other subnetworks at the site of T and other oral sites could also be determined, as shown in Supplementary Material 4.

**Table 6.**
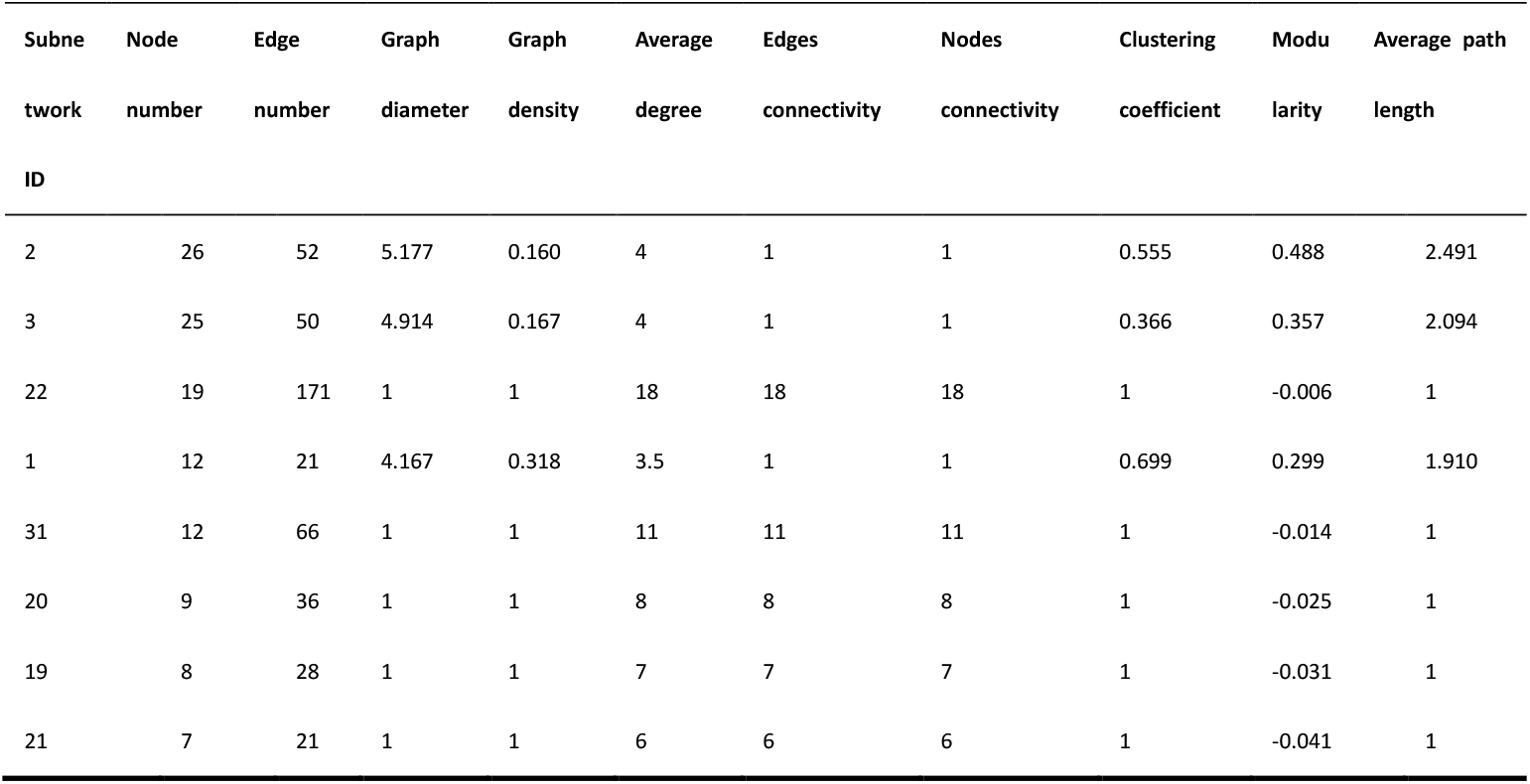
Topological properties of selected subnetworks.

| Subne | Node | Edge | Graph | Graph | Average | Edges | Nodes | Clustering | Modu | Average path |
| --- | --- | --- | --- | --- | --- | --- | --- | --- | --- | --- |
| work | number | number | diameter | density | degree | connectivity | connectivity | coefficient | larity | length |
| ID |  |  |  |  |  |  |  |  |  |  |
| 2 | 26 | 52 | 5.177 | 0.160 | 4 | 1 | 1 | 0.555 | 0.488 | 2.491 |
| 3 | 25 | 50 | 4.914 | 0.167 | 4 | 1 | 1 | 0.366 | 0.357 | 2.094 |
| 22 | 19 | 171 | 1 | 1 | 18 | 18 | 18 | 1 | -0.006 | 1 |
| 1 | 12 | 21 | 4.167 | 0.318 | 3.5 | 1 | 1 | 0.699 | 0.299 | 1.910 |
| 31 | 12 | 66 | 1 | 1 | 11 | 11 | 11 | 1 | -0.014 | 1 |
| 20 | 9 | 36 | 1 | 1 | 8 | 8 | 8 | 1 | -0.025 | 1 |
| 19 | 8 | 28 | 1 | 1 | 7 | 7 | 7 | 1 | -0.031 | 1 |
| 21 | 7 | 21 | 1 | 1 | 6 | 6 | 6 | 1 | -0.041 | 1 |

**Table 7.** Topological properties of selected nodes in subnetwork 2.

| Nodes | Degree | Weighted | Betweenness | Closeness | Eigenvector | RSR | Rank |
| --- | --- | --- | --- | --- | --- | --- | --- |
|  | Centrality | Degree | Centrality | Centrality | Centrality |  |  |
|  |  | Centrality |  |  |  |  |  |
| Treponema_bryantii | 9 | 7.815 | 119 | 0.023 | 1 | 3.111 | 1 |
| Desulfovibrio_piger | 7 | 6.067 | 155 | 0.025 | 0.517 | 2.822 | 2 |
| Maritalea_myrionectae | 7 | 6.163 | 53 | 0.022 | 0.638 | 2.747 | 3 |
| Flavonifractor_plautii | 7 | 6.164 | 19 | 0.022 | 0.638 | 2.712 | 4 |

#### 5. The microbial communities identified by MNetClass have high biological relevance to 5 oral sites

To enhance the credibility and interpretability of the experimental results, we will elucidate the reliability and biological significance of the key microbial subnetworks identified by MNetClass framework.

##### 5.1 Reliability evaluation of microbial community identification

We evaluated the reliability of the MNetClass framework in identifying key microbial communities across different oral sites, based on previous studies^70^. To enhance the robustness of our evaluation, we employed two assessment standards. The first standard utilized the expanded Human Oral Microbiome Database (eHOMD)^71^, which provides comprehensive data on microbial presence across various oral sites, including the core taxa for each site. The second approach involved a meta-analysis of the literature, using predefined keywords to identify core and “keystone” microbial communities from five oral sites, as reported in past studies^68,72,73^. These were summarized and used as a reference benchmark (details of the literature search methodology are provided in the Supplementary materials 4, with the benchmark summary in Supplementary Table 7). This approach is supported by recent reviews^74-76^ on oral microbiomes, which emphasize the significance of core and “keystone” taxa across different oral sites, an area of growing interest. Given the varying taxonomic levels (e.g., species, genus, or phylum) used in different studies, we mapped species and genus-level taxa to the phylum level using the Ribosomal Database Project47 (RDP version 11.5) to expand the scope of comparison. The microbial communities identified by different methods were also mapped to the phylum level for consistent comparison.

We compared the identified microbial communities with both benchmarks and calculated Precision, Recall, and F1-score metrics^77^. MNetClass was compared with five other community detection algorithms—edge betweenness^30^, label propagation^78^, leading eigenvector^79^, fast greedy^29^, Louvain^80^, as well as the manta clustering algorithm for weighted ecological networks^15^, and weighted gene co-expression network analysis (WGCNA)^49^. The results, shown in Figure 3(b) and Supplementary Table 6, indicate that, based on benchmark 1, MNetClass achieved the highest F1-score across all five oral sites, with the best performance in Precision and Recall in four sites. In benchmark 2, MNetClass achieved the highest F1-score in four oral sites, and the best performance in Precision and Recall in three sites. These results suggest that MNetClass outperforms the other seven algorithms by identifying a greater number of taxa that have been validated in databases or prior literature, making them more likely to play important physiological roles across different oral sites.

##### 5.2 Biological significance of identified central bacteria

*Treponema bryantii*, a rumen spirochete that interacts with cellulolytic bacteria^81,82^. *Maritalea myrionectae*, isolated from a culture of the marine ciliate Myrionecta rubra^83^. This suggests that these two central bacteria in subnetwork “2” of T may be associated with diet. *Desulfomonas pigra* is a type of sulfate-reducing bacteria (SRB)^84^. Emerging evidence indicates that SRB may play a significant role in human diseases, notably in relation to the clinical severity of human periodontitis^85^. *Flavonifractor plautii*, a gut probiotic^86^, has been documented to be involved in the metabolism of catechins in the human gastrointestinal tract, with its abundance positively correlated with green tea consumption^87^. Oral administration of *Flavonifractor plautii* has shown potential in alleviating inflammation in adipose tissue associated with obesity^88^, preventing arterial stiffness^89^, and exhibiting potential inhibitory effects on inflammatory bowel disease (IBD)^90^ and colorectal cancer^91^. The identification of central bacteria using MNetClass holds significant importance in understanding human health and interpreting physiological functions in different body sites. However, these central bacteria were not detected in the conventional differential analysis^92^ of microbial communities, as shown in Supplementary Material S5. This is because differential analysis is based on differences in the relative abundance of microbes in different body sites, and certain bacteria that play crucial roles may not exhibit significant differences in relative abundance across different sites. Consequently, this further underscores the contribution of MNetClass as a novel network analysis solution for microbiome studies.

### MNetClass equals or outperforms other algorithms on synthetic data sets

To evaluate the performance of MNetClass relative to other methods, we generated 50 synthetic datasets using FABIA^67^. Originally designed for evaluating biclustering of gene expression data, FABIA was selected for its ability to provide true labels, which serve as ground truth for performance assessment. Furthermore, it has been widely used in previous microbial network analysis studies^15^ to generate synthetic datasets for algorithm evaluation.

The accuracy of the identified microbial clusters was assessed using three metrics: complex-wise sensitivity (Sn), cluster-wise positive predictive value (PPV), and geometrical accuracy (Acc)^93^. Sn represents the proportion of correctly identified positive samples, reflecting how many of the true positives were accurately detected. PPV indicates the proportion of identified microbial clusters that are true positives. Acc, which combines Sn and PPV using their geometric mean, provides a balanced evaluation of both metrics. We compared MNetClass with five other community detection algorithms edge betweenness, label propagation, leading eigenvector, fast greedy, and Louvain as well as the biological weighted network clustering algorithm manta and weighted gene co-expression network analysis (WGCNA) using these evaluation metrics.

After partitioning the subnetworks using community detection methods, the most likely subnetworks to perform physiological functions were identified based on network topology analysis^60^. Previous studies typically used a single optimal topological property to extract subnetworks^94,95^; however, different topological properties yield different optimal subnetworks. As noted in recent literature, selecting subnetworks based on multiple topological properties is a more reasonable approach^96^. Therefore, we developed an integrated rank-sum ratio–entropy weight (RSR-EWM) evaluation model to provide a comprehensive score for network topological properties. We compared the performance of various community detection algorithms and the subnetworks selected based on single topological properties versus those selected using the RSR-EWM score across 50 synthetic datasets (Figure 4(a)). Except for the edge betweenness algorithm, where the subnetwork selected using a single topological property showed a higher ACC score than the RSR-EWM method, with no significant difference, all other algorithms performed better or equally well when combined with the RSR-EWM scoring. Notably, the Walktrap algorithm, when coupled with RSR-EWM, significantly outperformed the single topological property method across all three metrics. Furthermore, when generating box plots for these metrics, we excluded outliers and observed that MNetClass retained the highest number of points, indicating its stability and robustness across the 50 synthetic datasets.

**Figure 4.**
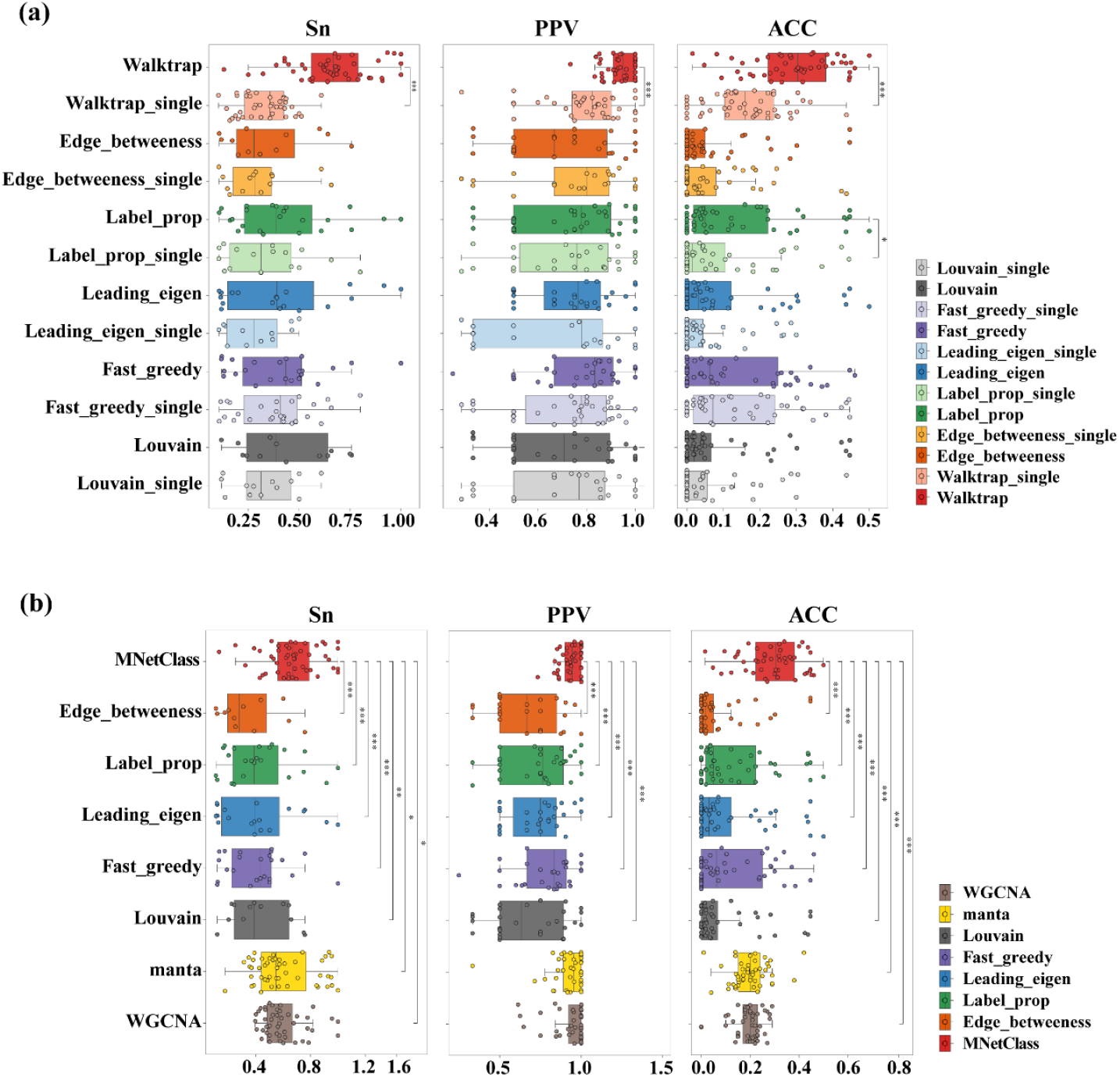
Performance of network clustering tools on 50 synthetic data sets generated with FABIA. (a) Evaluation of subnetwork selection using six methods, comparing the performance of individual topological properties versus the RSR-EWM scoring method, based on Sensitivity (Sn), Positive Predictive Value (PPV), and Accuracy (Acc). Sensitivity, PPV, Accuracy, and Separation (Sep) were calculated as previously described. “Single” in the legend denotes the use of a single topological property scoring method, while the absence of “single” indicates the use of the RSR-EWM scoring method by default. (b) Comparison of the microbial clusters identified by MNetClass and seven other algorithms, based on Sn, PPV, and ACC.

We compared MNetClass with seven other similar algorithms across 50 synthetic microbial datasets. For the microbial network construction step, edge betweenness, label propagation, leading eigenvector, fast greedy, Louvain, and the biological weighted network clustering algorithm manta did not specify a particular method for network construction, so we used the same approach as MNetClass. After network clustering, edge betweenness, label propagation, leading eigenvector, fast greedy, and Louvain selected key subnetworks based on the RSR-EWM topology scoring method, while manta selected the cluster with the highest proportion of strong nodes from the clustered groups as the key subnetwork. For WGCNA, originally designed for gene expression data, we adapted it for microbial co-occurrence network construction and module identification. We used default parameters to automatically construct the network and identify the modules most strongly associated with phenotype as the key subnetworks. The identified key subnetwork nodes were treated as predicted positives, and the Sn, PPV, and ACC were calculated by comparing these predicted positives with the true labels, as shown in Figure 3(b). MNetClass significantly outperformed the other six algorithms in Sn and ACC, and was significantly better than five other community detection algorithms in PPV, with no significant difference when compared to manta and WGCNA.

### Performance evaluation of MNetClass for identifying microbial key subnetwork and ASD prediction

#### 1. Identifying microbial key subnetworks across multiple cohorts

We applied our algorithm, MNetClass, to identify key microbial subnetworks across 10 independent ASD cohorts, following the workflow outlined in Figure 1. The identified microbial clusters are visualized in Figure 5(a). Compared to the original method, MNetClass revealed higher overlap in microbial clusters between the ASD and Healthy groups across all 10 cohorts, indicating greater consistency and reproducibility (Figures 5(f) and 5(g)). These results suggest that MNetClass effectively captures robust and shared microbial community structures.

**Figure 5.**
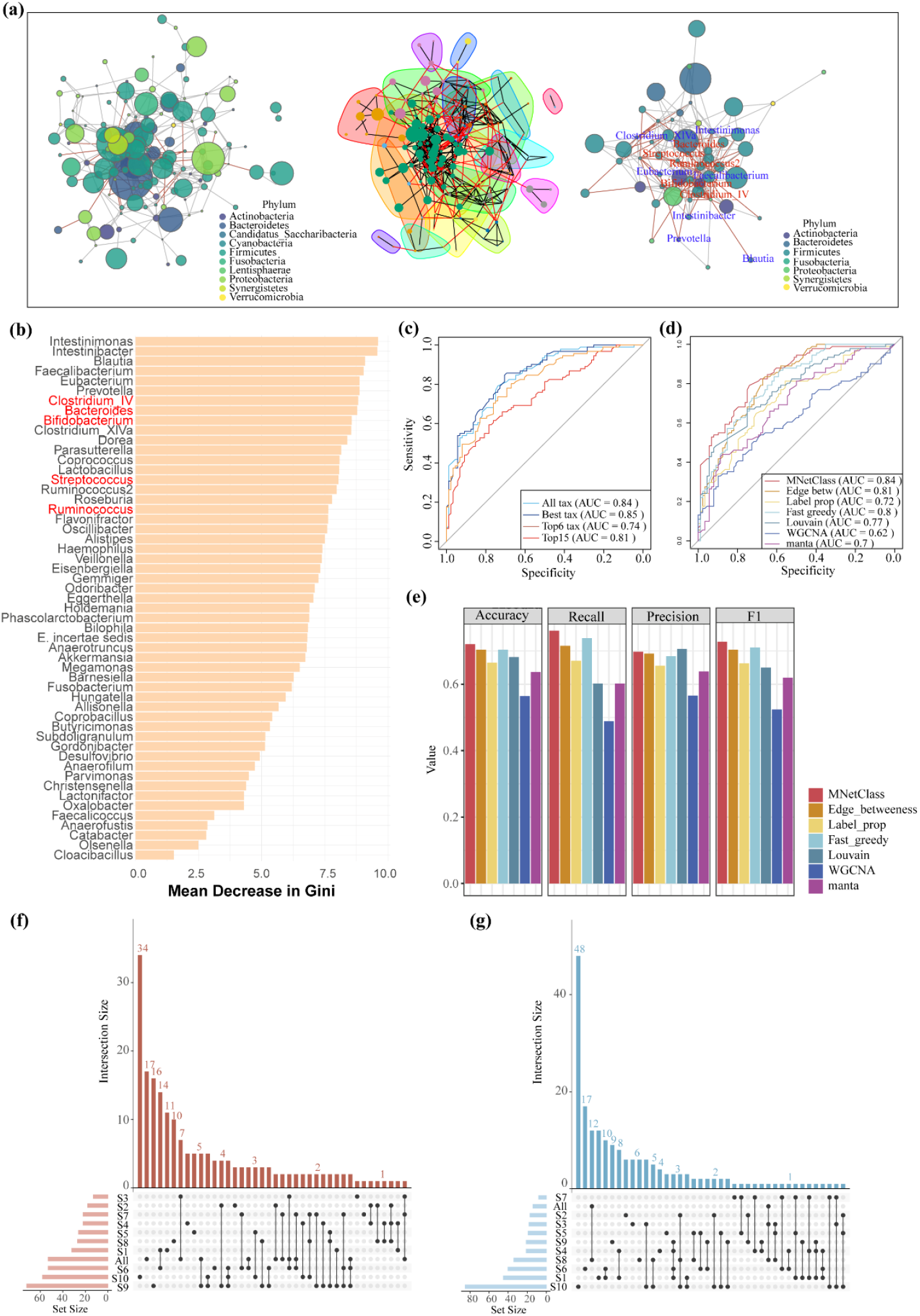
Benchmarking ASD prediction across multiple cohorts. (a) Visualization of the network construction, subnetwork partitioning, and subnetwork selection processes for ASD datasets using MNetClass. In the right panel, nodes with blue labels represent the top 10 genera ranked by importance in the random forest model shown in (b), while nodes with red labels represent genera previously validated as ASD-associated in the literature. (b) Importance ranking of microbial genera identified by MNetClass in the S1ASD cohort, based on the random forest model. (c) Receiver operating characteristic (ROC) curves for ASD prediction using the random forest model on cohorts S2 to S10. (d-e) ROC curves (d) and bar plots (e) comparing the Accuracy, Recall, Precision, and F1-score of random forest models constructed using microbial communities identified by MNetClass and seven other methods. (f-g) UpSet plots showing the microbial communities identified by MNetClass in the ASD and Healthy group across 10 cohorts.

#### 2. Predictive performance using identified subnetworks

To evaluate the predictive utility of the identified microbial key subnetworks, we utilized Project S1 as the training set and applied five-fold cross-validation to construct a random forest model. Feature selection was performed using recursive feature elimination (RFE) to determine the optimal number of features. Subsequently, we validated the model using external datasets, testing four configurations of features derived from the identified subnetworks: all taxa, the optimal number of taxa, and the top 6 and top 15 taxa ranked by feature importance (Figure 5(b)).

The ROC curves for these configurations are shown in Figure 5(c). Notably, using the optimal number of taxa, all taxa, and the top 15 taxa yielded superior predictive performance compared to the original method, with AUC scores of 0.85, 0.84, and 0.81, respectively. These results underscore the effectiveness of MNetClass in identifying predictive features and constructing robust classification models.

#### 3. Comparative performance with other algorithms

We compared the performance of microbial cluster identification by MNetClass with seven other algorithms for ASD prediction. Using Project S1 as the training set and the remaining datasets as validation sets, random forest models were constructed following the same pipeline. Five evaluation metrics were used: ROC-AUC, accuracy, recall, precision, and F1-score (Figures 5(d) and 5(e)).

MNetClass outperformed all other methods in ROC-AUC, accuracy, recall, and F1-score, while slightly underperforming Louvain in precision. These results demonstrate that MNetClass delivers superior performance in identifying meaningful microbial clusters for ASD prediction.

### Case study of MNetClass on oral microbial communities across age groups

#### 1. Subnetwork-related phenotypes as main drivers of sample grouping in oral microbiome data

We ran MNetClass on species-level oral microbiome data from all samples and performed a correlation analysis between the relative abundance of microbes in the identified key subnetwork and various phenotypes. Since the phenotypes-part, gender, and age group-are categorical variables, we conducted one-way ANOVA to assess whether the relative abundances of microbes significantly differed across categories (p ≤ 0.05). The results are shown in Figure 6(a) and in “Supplementary Table 8”

**Figure 6.**
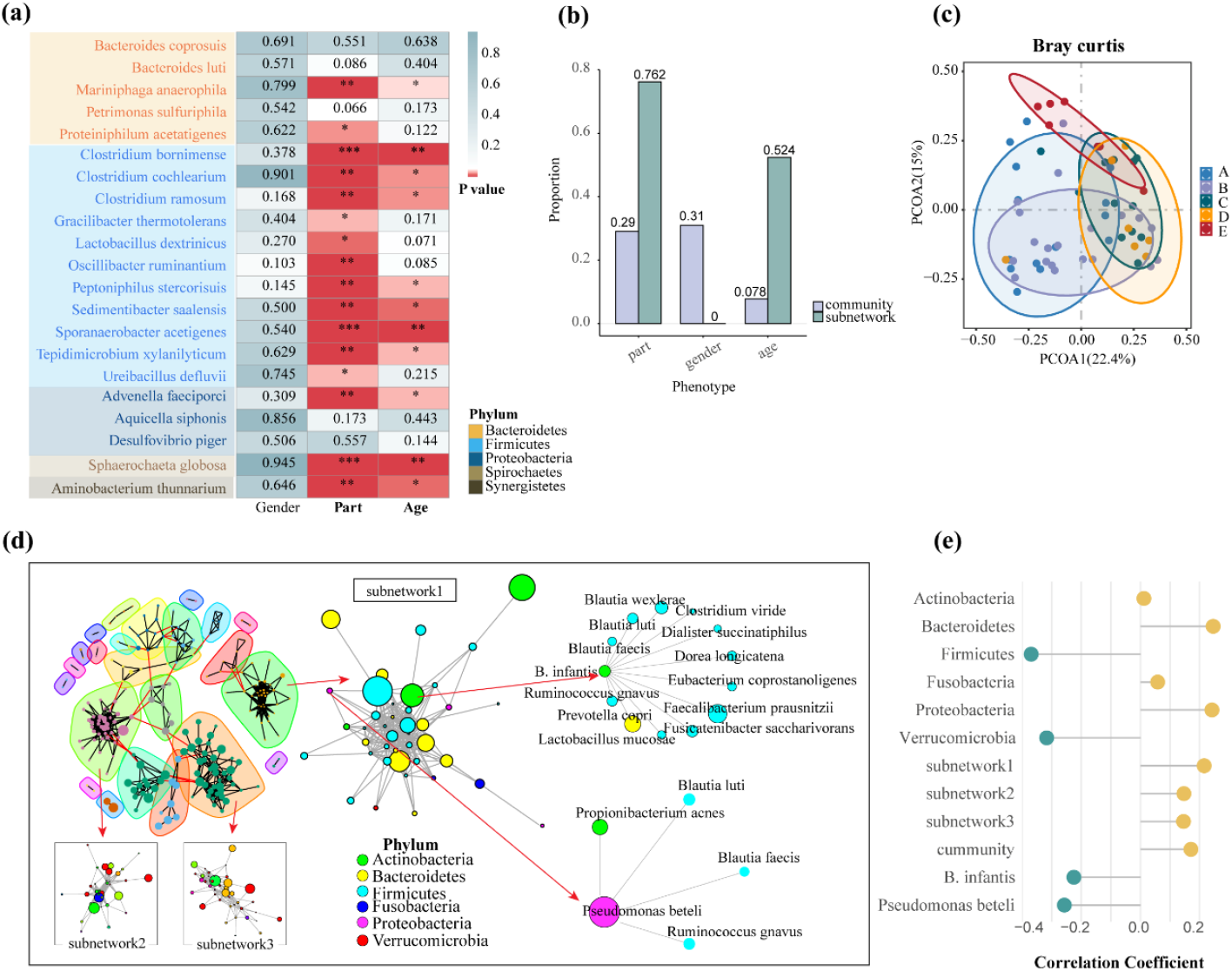
Application of MNetClass for indentifying age-associated oral bacteria. (a) Heatmap of p-values from ANOVA analysis comparing species in the key subnetworks identified by MNetClass across different phenotypes. In the heatmap, *p* ≤ 0.05 is marked with “*”, *p* ≤ 0.01 with “**”, *p* ≤ 0.001 with “***”, and *p* ≥ 0.05 with the corresponding p-value. (b) Proportions of species in the overall microbiome and those identified by MNetClass in the subnetworks that show significant correlation with different phenotypes, based on ANOVA analysis. Species with p ≤ 0.05 are considered significantly correlated. (c) Principal coordinate analysis (PCoA) of Bray-Curtis dissimilarities for sample compositions, with different colors representing different age groups: A (age 11–15), B (age 18–20), C (age 28–32), D (age 38–45), and E (age 50–65). (d) Visualization of network analysis results for GCF samples using MNetClass. Subnetworks 1–3 are ranked by the RSR-EWM scoring model, corresponding to rank-1, rank-2, and rank-3 subnetworks. (e) Average Spearman correlation coefficients between microbial relative abundances and age. “Subnetwork 1–3” represents the species within each subnetwork, while “Community” refers to the overall microbiome. Phyla identified by MNetClass, including *Actinobacteria, Bacteroidetes, Firmicutes, Fusobacteria, Proteobacteria*, and *Verrucomicrobia*, are also shown along with the species they contain.

The analysis revealed that the proportions of microbes with significant abundance differences across the phenotypes part, gender, and age in the identified subnetwork were 0.762, 0, and 0.524, respectively (Figure 6(b)). Additionally, we performed ANOVA on the relative abundance of all microbes with respect to the phenotypes, as detailed in “Supplementary Table 9” Here, the proportions of microbes with significant differences across the phenotypes were 0.290, 0.310, and 0.078 (Figure 6(b)).

These findings suggest that the subnetwork identified by MNetClass are more strongly correlated with the phenotype “part.” This is consistent with the original study’s findings, where Permutational Multivariate Analysis of Variance (PERMANOVA) indicated that “part” and “age” were the major drivers of sample classification. Furthermore, we performed PERMANOVA tests on the identified microbial community relative to different phenotypes. The p-values for “part,” “gender,” and “age” were 0.001, 0.886, and 0.001, respectively. These results demonstrate that our identified subnetwork enhances the significance of “part” and “age” as factors influencing groupings, while reducing the influence of “gender,” rendering it statistically insignificant. This confirms the need to study age-related oral microbiomes by separate anatomical sites, as suggested by the original analysis.

#### 2. MNetClass identifies age-related microbial communities across different oral sites

We used the GCF samples as an example for principal coordinates analysis (PCoA), as shown in Figure 6(c). A PERMANOVA test was performed to assess the influence of age on the sample grouping, with a significant p-value of 0.001. We then applied MNetClass to identify subnetworks (Figure 6(d)). Spearman correlation analysis was conducted between age and each species in the rank-1, rank-2, and rank-3 subnetworks, as well as in the time-varying microbial communities identified in the original study. The average correlation values are shown in Figure 6(e). The rank-1 subnetwork exhibited the highest average absolute Spearman correlation coefficient with age (0.2153, 0.1468, 0.1455, and 0.1705), indicating that the key subnetworks we selected are highly correlated with age. This may be due to the fact that, unlike in the original study, no predefined age groups were required, potentially reducing grouping-related biases and errors.

The phyla of the microbes identified in the key subnetworks by MNetClass include *Actinobacteria, Bacteroidetes, Firmicutes, Fusobacteria, Proteobacteria, and Verrucomicrobia*. In comparison, the phyla identified in the original study were *Actinobacteria, Bacteroidetes, Firmicutes, Proteobacteria, and Spirochaetes*, with only one microbial species not overlapping. This suggests that the microbial communities identified by MNetClass are consistent with those found in the original study.

Furthermore, we performed a topological analysis of the microbes in the identified subnetwork. Using RSR-EWM, we calculated the integrated topological scores of the nodes and selected those with the highest scores as the core microbes in the subnetwork. These microbes are more likely to play a physiological role in these subgroups. Notably, the central species identified in key subnetwork by MNetClass, *Bifidobacterium longum subsp. infantis* and *Pseudomonas beteli*, were also found in the original study’s key microbial species. Moreover, other species highly correlated with these two microbes may have significant biological relevance. For example, the increased abundance of *Dialister succinatiphilus* in periodontitis can serve as a biomarker, aiding in early diagnosis and monitoring disease progression^97^. *Lactobacillus mucosae*^*98,99*^ regulates the acidic environment of the oral cavity by producing lactic acid, inhibiting the growth of cariogenic bacteria and periodontal pathogens.

## DISCUSSION

Currently, most microbiome studies typically rely on statistical methods, such as the metagenomic wide association study (MWAS)^10^, and are not based on the principle of ecological theory^100^. Microbial network-based methods, such as NetShift^13^, NetMoss^14^ and manta^15^, are thus becoming powerful tools for elucidating the characteristics of microbiome ecosystems. However, most of these methods require a control group and are only suitable for measuring changes in microbial networks under healthy and diseased states. Moreover, very few methods exist for identifying subcommunities after clustering across ecological niches and defining the microorganisms that play an important role in these subpopulations. To address this issue, we established a control-independent microbial network clustering analysis framework, MNetClass, which expands the research methods on microbial interrelationships and ecological characteristics.

In MNetClass, we used the random walk community partition algorithm^18^ to divide the constructed association network into different subnetworks. This method is appropriate for the microbial network analysis constructed under different habitats because it does not require providing the number and size of partitions as parameters^19^. Additionally, the random walk algorithm can yield satisfactory results within a relatively small time complexity, ensuring the efficiency of our overall network analysis^56^. When evaluating the properties of subnetworks and their nodes, we use various topological property measurement indicators, such as network centrality. Nodes with the following properties can be discovered: a higher degree of connectivity to neighboring nodes, connections to a greater number of paths, closer proximity to other nodes in the network, or higher centrality among adjacent nodes. Therefore, with these indicators, we can identify the central species (or hubs)^28,66,101^ that may play important roles in maintaining the stability of the ecosystem. This step may garner significant interest among researchers because few methods focus on this topic.

To assess the topological properties of subnetworks and nodes of several different magnitudes, we used an integrated rank-sum ratio–entropy weight evaluation model^14^. This model mitigates the impacts of different indicators and different magnitudes on the comprehensive evaluation of multiple topological properties^102^. Furthermore, this model is an objective evaluation method that follows the principle that the greater the disparity in an indicator is, the lower its information entropy and the greater the amount of information it contains^63^. If all evaluation objects’ values for a particular indicator are equal, that indicator does not play a role in the comprehensive evaluation^103^. This method comprehensively scores and ranks the evaluation objects, facilitating the selection of dominant microbial communities at each site.

The focus of this study was to demonstrate an analytical workflow for microbial network clustering and central subcommunities identification, so the Spearman correlation coefficient, which is widely recognized as a classic measure for assessing the interactions between microbes, was employed to construct the microbial correlation network. Additionally, several other measurement methods, such as SparCC^45^, CCLasso^104^, and SPIEC-EASI^105^, can also be selected and utilized for network construction. Scientists can choose the appropriate method for measuring the interactions of microbes based on their research purpose. Our “MNetClass” R package will continue to be improved and updated in the future, such as by incorporating more interaction measurement options.

## ETHICS APPROVAL AND CONSENT TO PARTICIPATE

The studies involving human participants were reviewed and approved by the Medical Ethical Committee of School of Stomatology, Shandong University. The patients/participants provided their written informed consent to participate in this study.

## DATA AVAILABILITY

The raw sequence data reported in this paper have been deposited in the Genome Sequence Archive in National Genomics Data Center, China National Center for Bioinformation / Beijing Institute of Genomics, Chinese Academy of Sciences (GSA: CRA013444) that are publicly accessible at https://bigd.big.ac.cn/gsa/browse/CRA013444.

## SOFTWARE AVAILABILITY

The open-source MNetClass R package and tutorial are available at GitHub: https://github.com/YihuaWWW/MNetClass.

## AUTHORS’ CONTRIBUTIONS

All authors contributed to the pipeline development and workflow of analyses. The initial idea and framework were conceived by Qiang Feng and Yihua Wang. The pipeline was built and maintained by Yihua Wang, Qingzhen Hou and Bingqiang Liu. Yihua Wang, Qingzhen Hou and Fulan Wei contributed to the pipeline test and usage for network analyses. This protocol was written by Yihua Wang and Qiang Feng, and revised with from other authors’ suggestions.

## ACKNOWLEDGMENTS

We sincerely thank all colleagues in Shandong University-BOP Joint Oral Microbiome Laboratory for all of the kindly advices and support.

